# KSHV activates unfolded protein response sensors but suppresses downstream transcriptional responses to support lytic replication

**DOI:** 10.1101/442079

**Authors:** Benjamin P. Johnston, Craig McCormick

**Author notes:** Corresponding author: C.M.

## Abstract

Herpesviruses usurp host cell protein synthesis machinery to convert viral mRNAs into proteins, and the endoplasmic reticulum (ER) to ensure proper folding, post-translational modification and trafficking of secreted viral proteins. Overloading ER folding capacity activates the unfolded protein response (UPR), whereby displacement of the ER chaperone BiP activates UPR sensor proteins ATF6, PERK and IRE1 to initiate transcriptional responses to increase catabolic processes and ER folding capacity, while suppressing bulk protein synthesis. Kaposi’s sarcoma-associated herpesvirus (KSHV) can be reactivated from latency by chemical induction of ER stress, whereby the IRE1 endoribonuclease cleaves XBP1 mRNA, resulting in a ribosomal frameshift that yields the XBP1s transcription factor that transactivates the promoter of K-RTA, the viral lytic switch protein. By incorporating XBP1s responsive elements in the K-RTA promoter KSHV appears to have evolved a mechanism to respond to ER stress. Here, we report that following reactivation from latency, KSHV lytic replication causes activation of ATF6, PERK and IRE1 UPR sensor proteins. UPR sensor activation is required for efficient KSHV lytic replication; genetic or pharmacologic inhibition of each UPR sensor diminishes virion production. Despite strong UPR sensor activation during KSHV lytic replication, downstream UPR transcriptional responses were restricted; 1) ATF6 was cleaved to release the ATF6(N) transcription factor but known ATF6(N)-responsive genes were not transcribed; 2) PERK phosphorylated eIF2*α* but ATF4 did not accumulate as expected; 3) IRE1 caused XBP1 mRNA splicing, but XBP1s protein failed to accumulate and XBP1s-responsive genes were not transcribed. Remarkably, complementation of XBP1s deficiency during KSHV lytic replication by ectopic expression inhibited the production of infectious virions in a dose-dependent manner. Therefore, while XBP1s plays an important role in reactivation from latency, it inhibits later steps in lytic replication, which the virus overcomes by preventing its synthesis. Taken together, these findings suggest that KSHV hijacks UPR sensors to promote efficient viral replication while sustaining ER stress.

**Author summary:** Human herpesvirus-8 is the most recently discovered human herpesvirus, and it is the infectious cause of Kaposi’s sarcoma, which is why it’s also known as Kaposi’s sarcoma-associated herpesvirus (KSHV). Like all herpesviruses, KSHV replicates in the cell nucleus and uses host cell machinery to convert viral genes into proteins. Some of these proteins are synthesized, folded and modified in the endoplasmic reticulum (ER) and traverse the cellular secretory apparatus. Because the virus heavily utilizes the ER to make and process proteins, there is potential to overwhelm the system, which could impede viral replication and in extreme cases, kill the cell. Normally, when demands on the protein folding machinery are exceeded then misfolded proteins accumulate and activate the unfolded protein response (UPR). The UPR resolves ER stress by putting the brakes on synthesis of many proteins, while signaling to the nucleus to turn on a program that aims to correct this imbalance. Previous work has shown that KSHV is ‘wired’ to sense ER stress, which it uses to reactivate from a largely inactive state known as latency, in order to make more viruses. Specifically, a UPR sensor protein called IRE1 senses the accumulation of unfolded proteins in the ER and rededicates a gene called XBP1 to the production of a transcription factor called XBP1s through an unconventional cytoplasmic mRNA splicing event. XBP1s travels to the cell nucleus and stimulates the production of a collection of proteins that mitigate ER stress. In latently infected cells, XBP1s also binds to the KSHV genome and causes the production of K-RTA, a viral transcription factor that initiates the switch from latency to productive lytic replication. This achieves stress-induced initiation of KSHV replication, but nothing is known about how ER stress and the UPR affect progress through the KSHV replication cycle. Here we show that as KSHV replication progresses, all three known UPR sensor proteins, IRE1, ATF6 and PERK, are activated, which is required for efficient viral replication. Normally, activation of each of these three sensor proteins communicates a unique signal to the cell nucleus to stimulate the production of ER stress mitigating proteins, but in KSHV lytic replication all downstream communication is stymied. The failure to resolve ER stress would normally be expected to put the virus at a disadvantage, but we demonstrate that reversal of this scenario is worse; when we add extra XBP1s to the system to artificially stimulate the production of UPR responsive genes, virus replication is blocked at a late stage and no progeny viruses are released from infected cells. Taken together, these observations suggest that KSHV requires UPR sensor protein activation to replicate but has dramatically altered the outcome to prevent the synthesis of new UPR proteins and sustain stress in the ER compartment.

## Introduction

Proteins destined to be secreted are synthesized in the endoplasmic reticulum (ER), where they are folded by chaperone proteins and modified by glycosyltransferases and protein disulfide isomerases. Demands on the cellular protein folding machinery that exceed ER folding capacity trigger ER stress, which is the accumulation of ER-localized misfolded proteins [1]. This accrual of misfolded proteins activates the unfolded protein response (UPR) to mitigate the stress [2–4]. The UPR attempts to resolve ER stress by transiently attenuating translation, increasing the folding machinery, increasing lipid biogenesis to expand ER surface area, and degrading misfolded proteins in a process called ER-associated degradation (ERAD). If proteostasis is not re-established the UPR switches from an adaptive to an apoptotic response.

The UPR is coordinated by three transmembrane sensor proteins that sample the ER lumen; activated transcription factor-6 (ATF6), protein kinase R (PKR)-like endoplasmic reticulum kinase (PERK) and inositol-requiring enzyme 1 (IRE1). These sensor proteins are maintained in an inactive state by association of their lumenal domains with the ER chaperone BiP [5]. In response to ER stress, BiP is released to participate in re-folding reactions, allowing UPR sensor activation [6]. Together these three UPR sensors coordinate complementary aspects of an integrated transcription and translation program that mitigates ER stress.

ATF6 is a type II transmembrane protein and upon sensing unfolded proteins in the ER lumen, traffics to the Golgi apparatus, where it is cleaved by resident proteases site-1 protease (S1P) and site-2 protease (S2P) [7,8]. These cleavage events release the ATF6(N) fragment, which translocates to the nucleus to transactivate genes encoding chaperones, foldases and lipogenesis factors.

PERK activation by ER stress causes dimerization and autophosphorylation [9]. Active PERK phosphorylates serine 51 of eIF2α, which increases eIF2α affinity for its guanine exchange factor eIF2B [10,11]. This binding depletes the small pool of eIF2B, thereby inhibiting replenishment of the eIF2-GTP-Met-tRNA^Meti^ ternary complex required for translation initiation [12]. Bulk cap-dependent translation is attenuated, while a subset of uORF-containing mRNAs encoding stress response proteins are preferentially translated [13]. Activating transcription factor 4 (ATF4) is chief among these stress response proteins [14]; it translocates to the nucleus and drives the synthesis of genes products that mitigate ER stress by increasing a catabolic process known as autophagy and the antioxidant response [15,16]. ATF4 upregulates protein-phosphatase 1 α (PP1α) cofactor GADD34 (PPP1R15A) to resume translation by dephosphorylating eIF2α [17]. This direct control of protein synthesis by stress is known as the integrated stress response (ISR) [18,19]. During chronic or severe ER stress, ATF4 also upregulates the apoptotic effector CHOP [20–22].

IRE1 is a kinase and endoribonuclease ER transmembrane protein. ER stress triggers IRE1 dimerization, autophosphorylation, and activation of the cytosolic RNase domain that recognizes a conserved nucleotide sequence in two stem loops in the mRNA of *Xbp1* and excises a 26-nucleotide intron [23–25]. The tRNA ligase RTCB re-ligates cleaved *Xbp1* mRNA, generating a ribosomal frameshift that yields the active transcription factor XBP1s [26,27]. XBP1s translocates to the nucleus and drives production of chaperones, lipid synthesis proteins and proteins involved in ERAD [28]. The combined action of these gene products simultaneously increases ER folding capacity and decreases the load via ERAD. In a process called regulated IRE1-dependent decay (RIDD), IRE1 can cleave ER-targeted mRNAs with a stem loop that resembles that of XBP1, triggering mRNA degradation, which like PERK, is thought to attenuate translation [29,30].

UPR signalling is involved in many normal physiological processes and dysregulation can contribute to the development and progression of human disease. The UPR is important for differentiation of professional secretory cells like pancreatic beta cells and antibody-secreting plasma B cells [31–33], and it is mechanistically linked to diseases such as Crohn’s disease and type 2 diabetes [34,35]. Several viruses have been shown to modulate UPR signalling during infection [36].

KSHV is a gammaherpesvirus that causes Kaposi’s sarcoma (KS), primary effusion lymphoma (PEL) and multicentric Castleman’s disease (MCD) [37–39]. Like all herpesviruses, KSHV can establish a quiescent form of infection known as latency in which viral gene expression is severely restricted and the genome is maintained as a nuclear episome. The virus can latently infect a variety of cell types, but it is thought that life-long infection of human hosts is primarily enabled by latent infection of immature B lymphocytes [40], followed by viral reprogramming into an intermediate cell type that resembles a plasma cell precursor [41–43].

An essential feature of latency is reversibility, which is required for viral replication and production of viral progeny. The true physiologic cues for KSHV lytic reactivation are not known, but *in vitro* studies have implicated ER stress-activated signalling pathways. Reactivation from latency requires the immediate-early lytic switch protein <u>r</u>eplication and <u>t</u>ranscriptional <u>a</u>ctivator (RTA), a transcription factor that initiates a temporal cascade of gene expression [44,45]. RTA recruits host cell co-factors to transactivate viral early gene promoters required for genome replication [46]. KSHV has usurped the IRE1/XBP1s pathway to regulate latent/lytic switch. XBP1s binds to canonical response elements in the RTA promoter and drives synthesis of the RTA lytic switch protein required for lytic reactivation [47–49]. Functional XBP1s binding motifs have also been found in the promoter for v-IL6 [50], the KSHV homolog of human IL6, which plays key roles in PEL and MCD cancers [51–53]. Thus, UPR activation causes reactivation from latency and initiation of lytic gene expression. Interestingly, because the XBP1 transcription factor is required for normal B cell differentiation into mature plasma cells [32,33], it is likely that the viral acquisition of XBP1s target sequences hardwires KSHV reactivation from latency to terminal B cell differentiation [47], although the physiological consequences of this link are not known. In sharp contrast to these mechanistic linkages, nothing is known about how ATF6 and PERK impact latent KSHV infection or reactivation from latency.

KSHV replicates in the nucleus and usurps host cell machinery to transcribe, post-transcriptionally modify, export and translate viral mRNAs. KSHV encodes structural and non-structural proteins that are folded in the ER and traverse the secretory apparatus. UPR activation during herpesvirus lytic replication has been reported, and there is evidence for UPR sensor engagement by specific gene products, rather than simply by exceeding ER folding capacity [54–56]. HSV-1 gB glycoprotein can bind PERK to promote virus protein production [57]. Nothing is known about how KSHV lytic gene products affect UPR.

To better understand how KSHV usurps the UPR, we investigated UPR activation following reactivation from latency in B cell- and epithelial cell-based models. We report that all three proximal UPR sensors are activated following reactivation from latency; ATF6 is cleaved, PERK and eIF2α are phosphorylated, and IRE1 is phosphorylated and *Xbp1* mRNA is spliced. Furthermore, we determined that UPR sensor activation is pro-viral; pharmacologic or genetic inhibition of each UPR sensor diminished virion yield from infected cells. Surprisingly, viral proteins accumulated despite sustained phosphorylation of eIF2α throughout the lytic cycle, suggesting that viral messenger ribonucleoproteins (mRNPs) may have unique properties that ensure priority access to translation machinery during stress. Remarkably, activation of proximal UPR sensors during lytic replication failed to elicit any of the expected downstream effects: ATF6(N) and XBP1-targets genes were not upregulated and ATF4 was not translated. This suggests that the virus requires proximal activation of UPR sensors, but prevents the UPR from mitigating stress and restoring ER homeostasis. Indeed, despite clear evidence of IRE1 activation and XBP1 mRNA splicing, XBP1s protein failed to accumulate during KSHV lytic replication. Remarkably, complementation of XBP1s deficiency during KSHV lytic replication by ectopic expression inhibited the production of infectious virions. Therefore, while XBP1s plays an important role in reactivation from latency, it inhibits later steps in lytic replication, which the virus overcomes by tempering its synthesis. Taken together, these findings suggest that KSHV hijacks UPR sensors to promote efficient viral replication instead of resolving ER stress.

## Results

### KSHV lytic replication activates IRE1 and PERK but downstream UPR transcription factors XBP1s and ATF4 fail to accumulate

ER stress can induce KSHV lytic replication via IRE1 activation and downstream XBP1s transactivation of the latent/lytic switch protein RTA [47,48] (Supplemental Fig. 1A and B). However, it is not known whether the burden of synthesizing secreted KSHV proteins causes ER stress over the course of the lytic replication cycle. To test UPR activation we used the TREx BCBL1-RTA cell line that expresses RTA from a doxycycline (dox)-regulated promoter to reactivate KSHV from latency [58]. We treated cells with dox for 0, 24, and 48 hours (h) and immunoblotted for UPR proteins to determine their activation state and KSHV proteins to monitor the progression of the lytic cycle from early (ORF45) to late (ORF65) stages (Fig. 1A). As a positive control to ensure that UPR sensors were intact in our system, we also treated latent and lytic cells with the SERCA (sarco/endoplasmic reticulum Ca^2+^-ATPase) inhibitor thapsigargin (Tg) to pharmacologically induce ER stress [59]. After 2 h of Tg treatment of latently-infected cells, IRE1 α was phosphorylated as determined by a migration shift in a total protein blot, and a semiquantitative *Xbp1* RT-PCR splicing assay revealed that the majority of *Xbp1* mRNA was spliced, causing a ribosomal frameshift and synthesis of XBP1s protein. Tg activated the ISR as determined by phosphorylation of PERK (also determined by a migration shift in total protein) and its downstream target eIF2α-Ser51, which resulted in translation of the uORF-containing mRNA ATF4. Lysates collected at 24 and 48 h post dox addition displayed strong accumulation of ORF45 and ORF65, respectively, and activation of both IRE1α and PERK pathways. However, PERK activation and eIF2α phosphorylation did not result in full ISR activation in lytic cells because there was minimal induction of ATF4. ATF4 also failed to accumulate in Tg-treated lytic cells, suggesting that lytic replication prevents downstream execution of the ISR when eIF2α is phosphorylated. Tg-treated lytic cells also displayed reduced total IRE1α and XBP1s levels, even though *Xbp1* mRNA splicing was similar to Tg-treated latent cells. These data demonstrate that KSHV lytic replication activates IRE1α and PERK UPR sensor proteins but prevents the accumulation of XBP1s and ATF4 transcription factors required for downstream transcriptional responses.

**Fig. 1:**
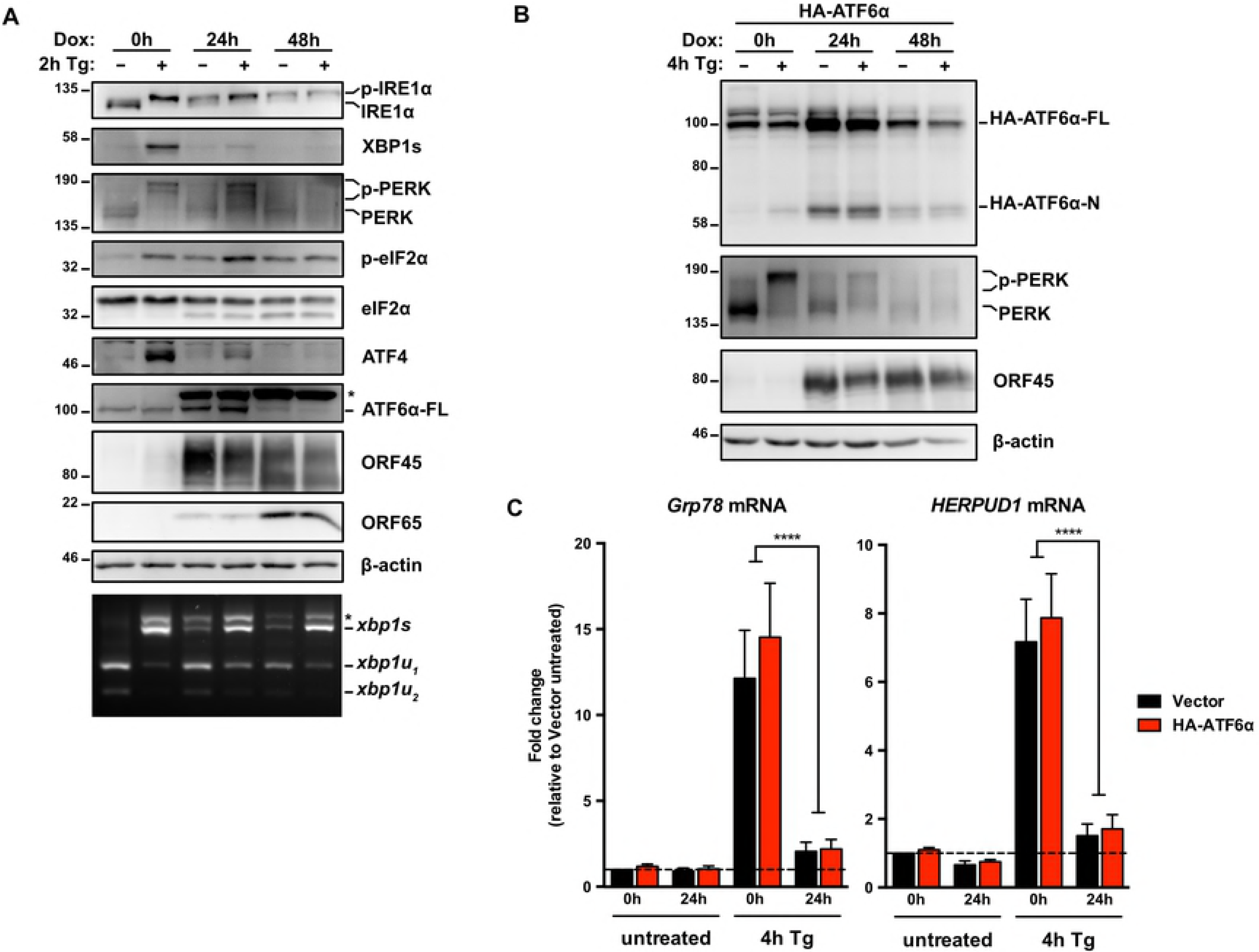
KSHV lytic replication activates IRE1, PERK and ATF6 but downstream UPR transcription factors are inhibited. (A) TREx BCBL1-RTA cells were treated with 1μg/ml doxycycline (dox) for 0, 24 and 48 hours to induce lytic replication followed by 75nM thapsigargin (Tg) for 2 hours prior to harvesting for protein and total RNA. Whole cell lysates were analyzed by immunoblots for UPR markers (IRE1α, XBP1s, PERK, phospho- and total eIF2α, ATF4, and full length ATF6α. Migration shift in PERK and IRE1α immunoblots correspond to phosphorylation. KSHV proteins ORF45 and ORF65 were probed for to indicate induction of early lytic and late lytic, respectively. β-actin was used as a loading control. The asterisk (*) corresponds to an unknown protein species that cross-reacted with ATF6 anti-sera. (Bottom) XBP1 mRNA was amplified by RT-PCR, digested with PstI (cleaves unspliced XBP1 isoform only), and separated on SYBR Safe-stained agarose gel. The asterisk (*) corresponds to xbp1u-xbp1s hybrid cDNA. Representative immunoblot and agarose gel of two independent experiments are shown (B) TREx BCBL1-RTA cells were transduced with lentiviral expression vector encoding HA-ATF6α and selected for with 1μg/ml puromycin. Following selection, cells were treated with 1μg/ml doxycycline for 0, 24 and 48 hours and treated with 75nM thapsigargin for 4 hours prior to harvest. Whole cell lysates were analyzed by immunoblots for HA epitope tag, PERK, ORF45 and β-actin (loading control). Immunoblot shown is representative of two independent experiments. (C) As in (B), HA-ATF6a-transduced TREx BCBL1-RTA were treated with doxycycline for 0 and 24 hours and then treated with 75nM thapsigargin for 4 hours prior to total RNA isolation. mRNA levels of ATF6α target genes GRP78 and HERPUD1 were measured by qPCR. Changes in mRNA levels were calculated by the ΔΔ*Cq* method and normalized using 18S rRNA as a reference gene. An average of 3 independent experiments are graphed and error bars denote SEM. Two-way ANOVA and a post-hoc multiple comparisons test were done to determine statistical significance (****, *p* value < 0.0001).

### KSHV lytic replication activates ATF6 but downstream UPR transcriptional responses are inhibited

The third UPR sensor ATF6 is cleaved in the Golgi during ER stress to release its active N-terminal fragment ATF6(N) that translocates to the nucleus and transactivates a variety of UPR genes [8]. We observed elevated total endogenous ATF6 levels at 24 h post dox addition, which diminished over the following 24 h (Fig. 1A). However, the ATF6 antibody that we employed in this study could not detect endogenous ATF6(N) in any samples, including Tg-treated positive controls. Previous reports show that the N-terminal fragment is susceptible to degradation [8,60]. Interestingly, we detected the emergence of an additional high MW species that reacted with the ATF6 antiserum in lysates from dox-treated cells (Fig. 1A, marked with a *), which to our knowledge has not been reported in the literature before.

To further investigate ATF6 cleavage in this system, we transduced TREx BCBL1-RTA cells with a lentiviral vector encoding HA-epitope-tagged full-length ATF6 (HA-ATF6αFL) [61]. Transduced cells treated with Tg revealed the detection of the N-terminal fragment (Fig. 1B, lane 2). Consistent with endogenous ATF6 in lytic cells (Fig. 1A), we observed the accumulation of full-length HA-ATF6 at 24 hours post-dox followed by return to basal levels by 48 hours post-dox. This ectopic expression system allowed us to monitor cleavage of the ~100 kDa HA-ATF6α FL precursor into the ~60 kDa HA-ATF6α(N) fragment during lytic replication (Fig. 1B, lanes 3-6). Indeed, by 24 hours post-dox the levels of HA-ATF6α(N) were higher than latently infected cells treated with Tg. Interestingly, ATF6-target genes, GRP78 and HERPUD1 [62–64], were not transactivated during lytic replication, even in cells treated with Tg (Fig. 1C). These data show that ATF6 is activated during lytic replication but the N-terminal transcription factor is unable to transactivate canonical target genes.

### PERK-dependent eIF2α phosphorylation during the KSHV lytic cycle

Because eIF2α can be phosphorylated on serine 51 by four possible eIF2α kinases activated by different stresses (ER stress activates PERK, dsRNA activates PKR, nutrient stress activates GCN2, oxidative stress activates HRI [9,65–67]), we investigated the specific contribution of PERK to eIF2α phosphorylation during KSHV lytic replication. TREx BCBL1-RTA cells were treated with PERK inhibitor GSK2606414 (PERKi) [68,69] concurrent with the addition of dox. After 24 h and 48 h of treatment PERKi caused a reduction in PERK and eIF2α phosphorylation compared to cells treated with dox alone (Fig. 2). Importantly, PERKi had no appreciable effect on IRE1α activation, indicating that ER stress is still manifested during KSHV lytic replication when PERK is inhibited, and PERK inhibition does not heighten this response. Interestingly, the level of eIF2α phosphorylation following PERKi treatment did not return to baseline levels observed in latently-infected TREx BCBL1-RTA cells, suggesting that other eIF2α kinases besides PERK may be activated during lytic replication. Furthermore, treating lytic cells with PERKi revealed that even though ATF4 fails to accumulate during lytic replication with or without Tg treatment, PERK-mediated phosphorylation of eIF2α may still inhibit bulk protein synthesis, as evidenced by the increased levels of the early viral gene product ORF45 (Fig. 2A, compare lanes 9 vs 10 and lanes 11 vs 12).

**Fig. 2:**
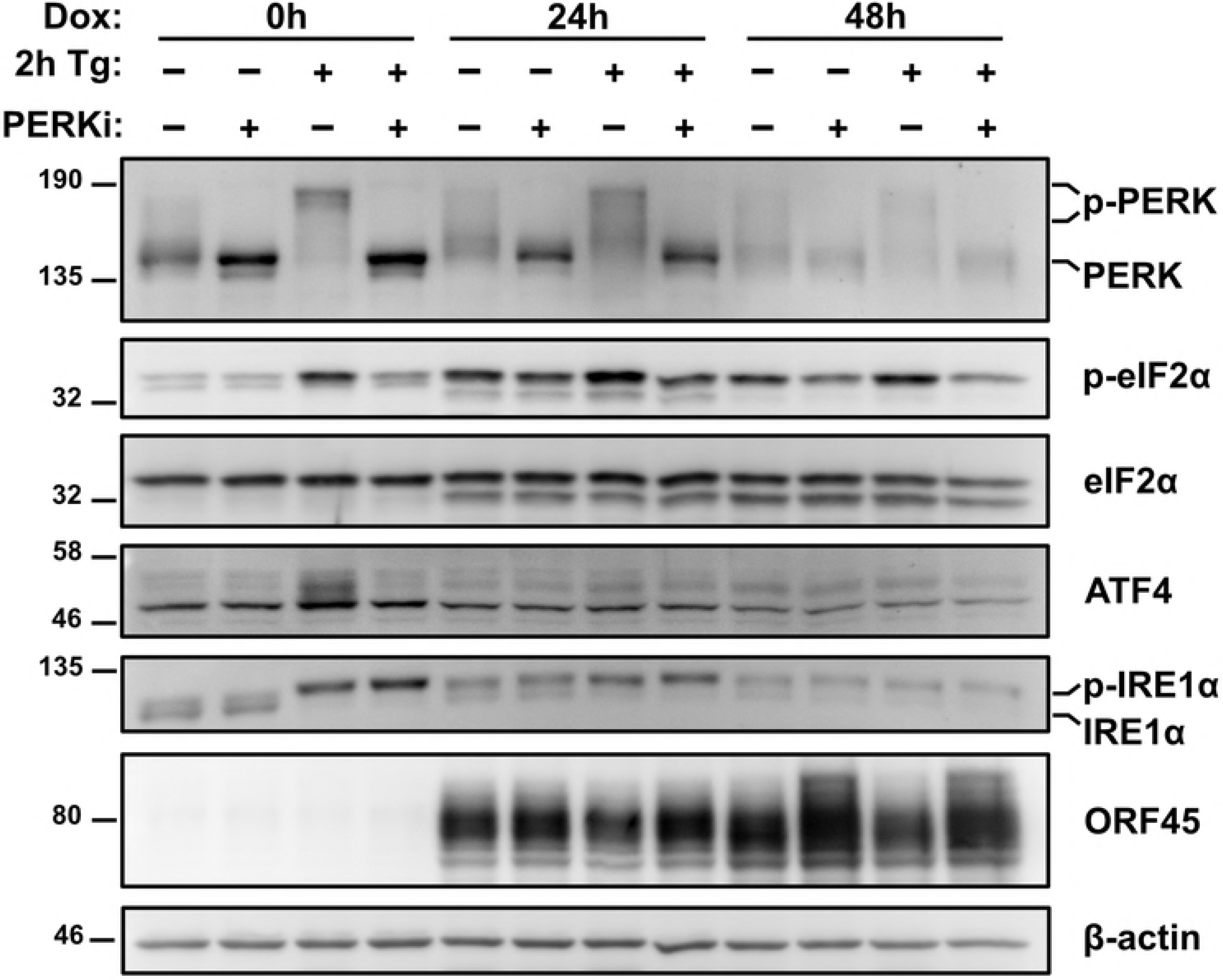
PERK-dependent eIF2α phosphorylation during the KSHV lytic cycle. TREx BCBL1-RTA cells were treated with 1pg/ml doxycycline −/+ 500 nM of the PERK inhibitor GSK2606414 (PERKi) for 0, 24 and 48 hours and treated with or without 75 nM thapsigargin for 2 hours prior to harvest. Whole cell lysates were analyzed by immunoblots for PERK, phospho and total eIF2α, ATF4, IRE1α, and ORF45. Migration shift in PERK and IRE1α immunoblots correspond to phosphorylation. β-actin was used as a loading control. Immunoblots shown are representative of two independent experiments.

### Failure of XBP1s and ATF4 to accumulate is not specific to the type or duration of ER stress

We previously observed UPR sensor activation but failure to accumulate active UPR transcription factors XBP1s, ATF4 and ATF6(N) following a brief 2 h pulse of Tg (Figs. 1A, B). We confirmed these observations by treating latent and lytic TREx BCBL1-RTA cells with Tg or tunicamycin (Tm), which induces ER stress by blocking protein N-linked glycosylation in the ER [70,71], over a range of incubation times (1h, 4h, 8h). Maximal *Xbp1* mRNA splicing was observed after latent cells were Tg-treated for 1 h or Tm-treated for 4 h (Fig. 3); these slower kinetics are consistent with known properties of Tm, which relies on new protein synthesis to trigger ER stress [72]. As expected, in latently infected cells, *Xbp1* splicing and XBP1s and ATF4 protein accumulation were observed throughout the 8 h course of Tg or Tm treatment. By contrast, during KSHV lytic replication, regardless of duration of Tg or Tm treatment, comparable levels of spliced *Xbp1* mRNA and phospho-eIF2α were detected but XBP1s and ATF4 proteins failed to accumulate. Consistent with previous results, there were slightly higher levels of unspliced *Xbp1* mRNA observed during the lytic cycle that correlated with reduced total IRE1α protein levels, but this was only observed in Tg-treated cells; Tm treatment converted the bulk of the *Xbp1* mRNA pool to the spliced form by 4 h post-treatment, even though IRE1α accumulation was still blocked. Consistent with our previous observations total ATF6 levels increased during the lytic cycle, and protein species with different electrophoretic mobilities could be detected by the ATF6 antiserum. Extended Tg or Tm treatments caused accumulation of the ATF6(N) transcription factor exclusively in the lytic cells. Taken together, these observations confirm that XBP1s and ATF4 transcription factors fail to accumulate in the KSHV lytic cycle despite robust activation of UPR sensors after chemically-induced ER stress.

**Fig. 3:**
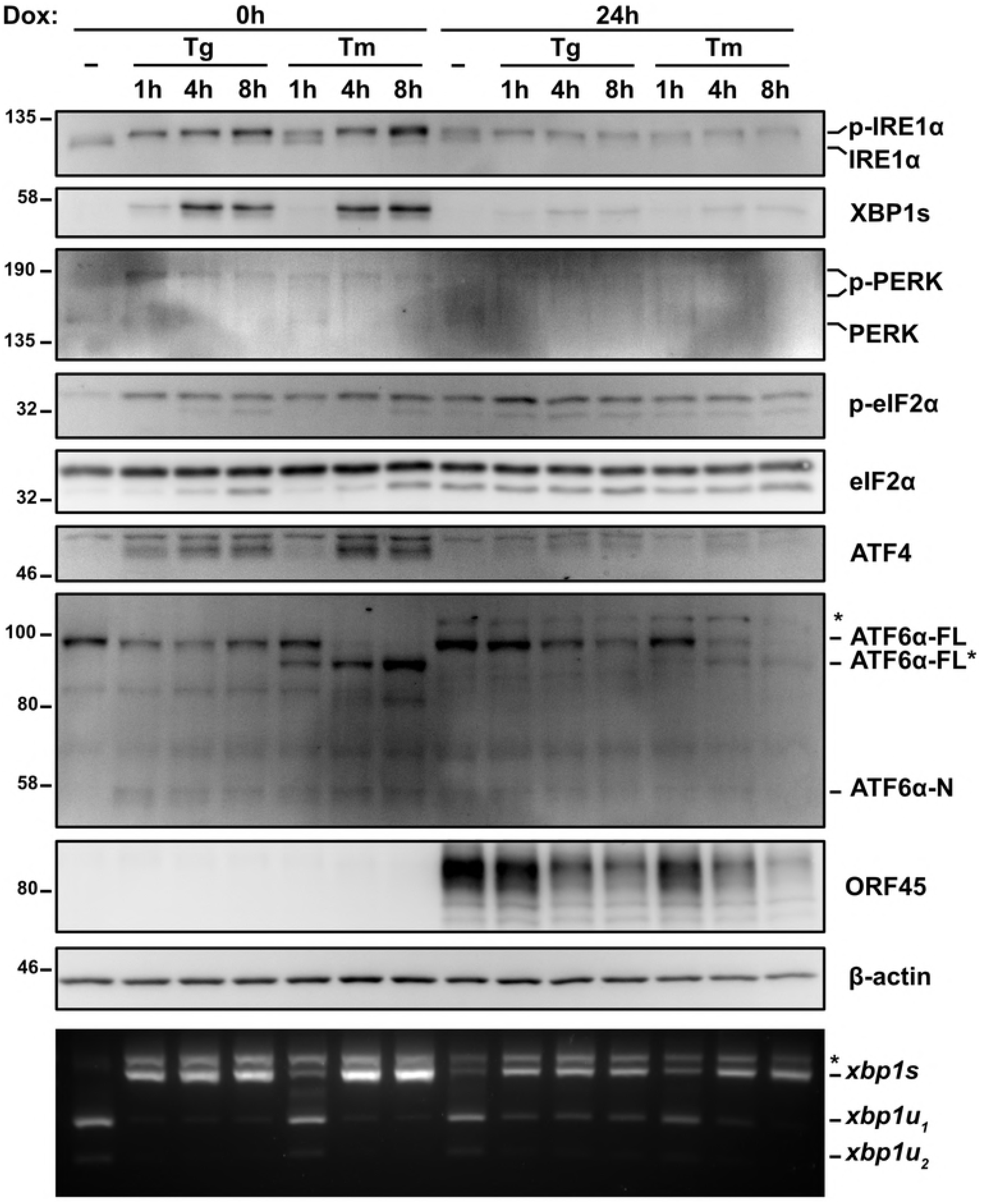
The type or duration of ER stress does not account for the lack of ATF4 and XBP1s. TREx BCBL1-RTA cells were pretreated with doxycycline for 0 or 24 hours and treated with 75nM Tg or 5μ/ml tunicamycin (Tm) for 1, 4, or 8 hours prior to harvest for either protein or RNA. Whole cell lysates were analyzed by immunoblots for UPR markers (IRE1α, XBP1s, PERK, phospho- and total eIF2α, ATF4, and ATF6α). Migration shift in PERK and IRE1α immunoblots correspond to phosphorylation. KSHV protein ORF45 was probed to show lytic replication and β-actin was used as a loading control. (Bottom) XBP1 RT-PCR splicing assay was performed as previously indicated. (*) corresponds to xbp1u-xbp1s hybrid cDNA. Immunoblots and agarose gels are representative of two independent experiments.

### XBP1s target genes are not transactivated by XBP1s during lytic replication

To better understand the kinetics of IRE1 activity during KSHV lytic replication we reactivated TREx BCBL1-RTA cells from latency using dox and monitored IRE1 activity over a 24 h time course. Spliced XBP1 mRNA and XBP1s protein were detected by 6 h post-dox addition, concomitant with a modest increase in IRE1 phosphorylation as determined by retarded IRE1 mobility on an immunoblot probed with an anti-IRE1α antibody (Fig. 4A). This induction of *Xbp1* splicing corresponds with the increase in the early viral protein ORF45. By 18 hours post dox addition, which coincides with the beginning of viral genome replication in this model [58], IRE1 phosphorylation, *Xbp1* splicing and XBP1s accumulation had peaked, with almost 40% of *Xbp1* mRNA in the spliced isoform; however, these phenotypes were surpassed by the Tg positive control (Fig. 4A and B). We wanted to test if the low levels of XBP1s that we observe during lytic replication would be sufficient to induce synthesis of XBP1s target genes. We harvested RNA from latent and lytic (24 h post dox addition) TREx BCBL1-RTA cells treated with either Tg or Tm or left untreated and performed semiquantitative RT-PCR analysis. As previously observed, *Xbp1* mRNA was efficiently spliced following Tg or Tm treatment, both in latent and lytic samples, while moderate splicing was observed in the mock-treated lytic samples (Fig. 4C). RT-qPCR was performed to measure relative levels of canonical XBP1s target genes EDEM1 and ERdj4 [73] (Fig. 4D). Indeed, Tg and Tm treatments caused dramatic accumulation of EDEM1 and ERdj4 transcripts in latently infected cells, whereas these transcripts remained at low levels in lytic cells treated with Tg, Tm or mock-treated. Thus, low expression of XBP1s during lytic replication did not induce synthesis of canonical XBP1s target gene products involved in ER stress mitigation.

**Fig. 4:**
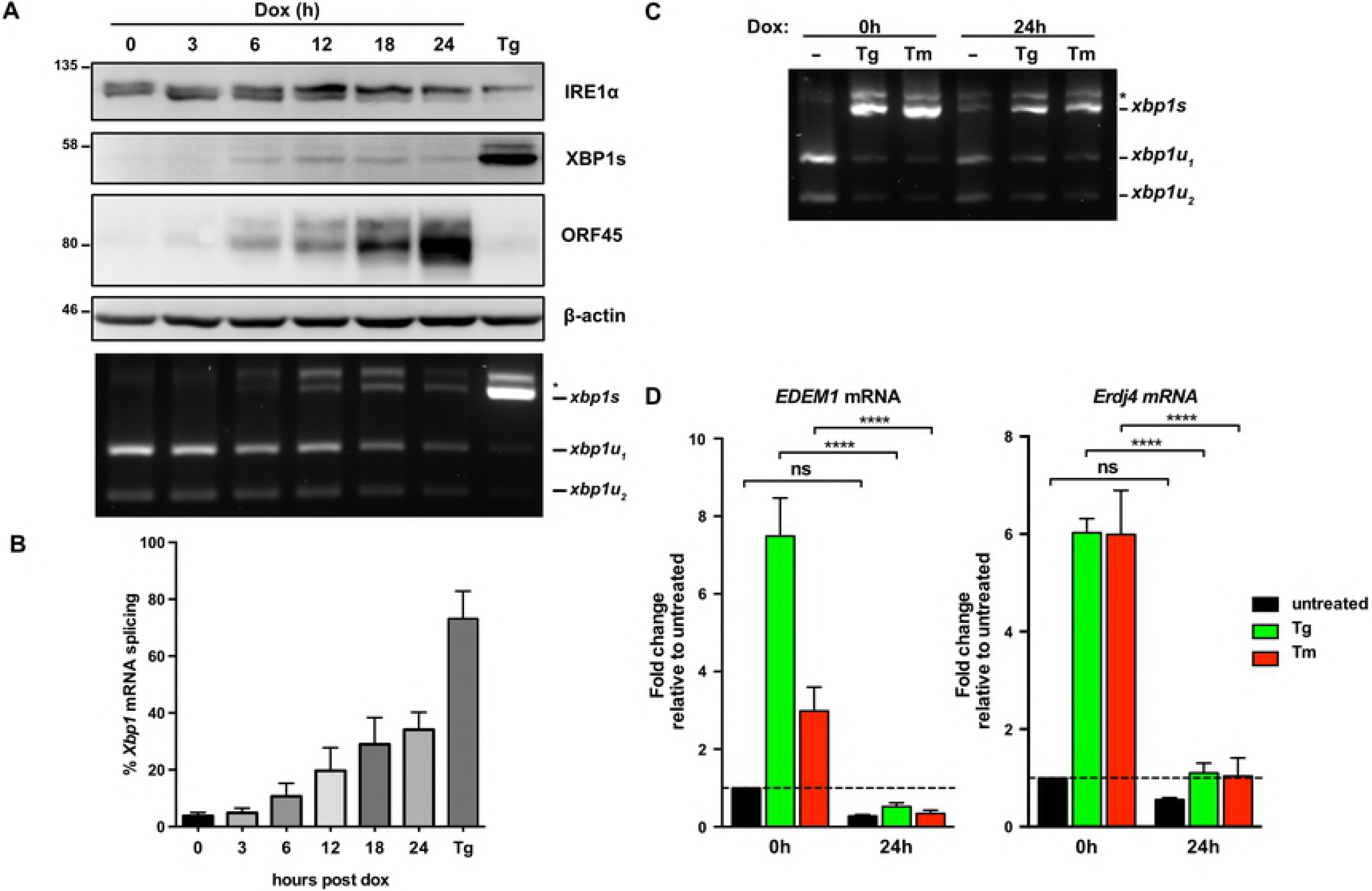
Low levels of XBP1s induced in lytic are insufficient to upregulate XBP1s-target genes. (A) TREx BCBL1-RTA cells were treated with 1μg/ml doxycycline (dox) for the indicated times or treated with 75nM Tg for 4 hours (as a positive control) and harvested for either protein or RNA. Whole cell lysates were analyzed by immunoblots for IRE1α, XBP1s, ORF45 and β-actin (loading control). (Bottom) XBP1 RT-PCR splicing assay was performed as previously indicated. (*) corresponds to xbp1u-xbp1s hybrid cDNA. A representative immunoblot and agarose gel for three independent experiments are shown. (B) Densitometry analysis from semi-quantitative XBP1 RT-PCR splicing assay in (A) was used to calculate the percentage of XBP1 in the spliced isoform. The mean of three independent experiments are graphed and error bars represent the standard deviation of the mean. (C) TREx BCBL1-RTA cells were treated with 1μg/ml doxycycline (dox) for 0 or 24 hours and treated with 75nM Tg or 5μg/ml Tm for 4 hours prior to RNA isolation. XBP1 mRNA splicing was determined by semi-quantitative RT-PCR splicing assay as previously described. The gel shown is representative of two independent experiments. (D) Total RNA samples from (C) were used to measure the mRNA levels of XBP1 target genes EDEM1 and ERdj4 by qPCR. Changes in mRNA levels were calculated by the ΔΔ*Cq* method and normalized using 18S rRNA as a reference gene. An average of 4 independent experiments are graphed and error bars denote SEM. Two-way ANOVA and a post-hoc multiple comparisons test were done to determine statistical significance (****, *p* value < 0.0001).

### UPR sensor activation supports efficient KSHV replication

We next used a combination of genetic and pharmacologic approaches to inhibit each of the UPR sensor proteins and measure their impact on KSHV lytic replication. To investigate the role of ATF6, we silenced ATF6α expression in TRex BCBL-RTA cells with shRNAs (Fig. 5A) and collected cell supernatants from ATF6-silenced or control cells at 48 h post-dox addition. Cell supernatants were processed to measure relative levels of released capsid-protected viral genomes by qPCR. ATF6 knockdown reduced virus titre by ~50% compared to cells transduced with non-targeting shRNA (Fig. 5B). To determine whether activation of PERK or downstream engagement of the ISR are important for viral replication TREx BCBL1-RTA cells were treated with the selective PERK inhibitor GSK2606414 (PERKi) or the ISR inhibitor ISRIB, a small molecule that blocks phospho-eIF2α-mediated inhibition of translation by maintaining active eIF2B [74,75]. PERKi and ISRIB each inhibited viral particle release by ~50% (Fig. 5C).

**Fig. 5:**
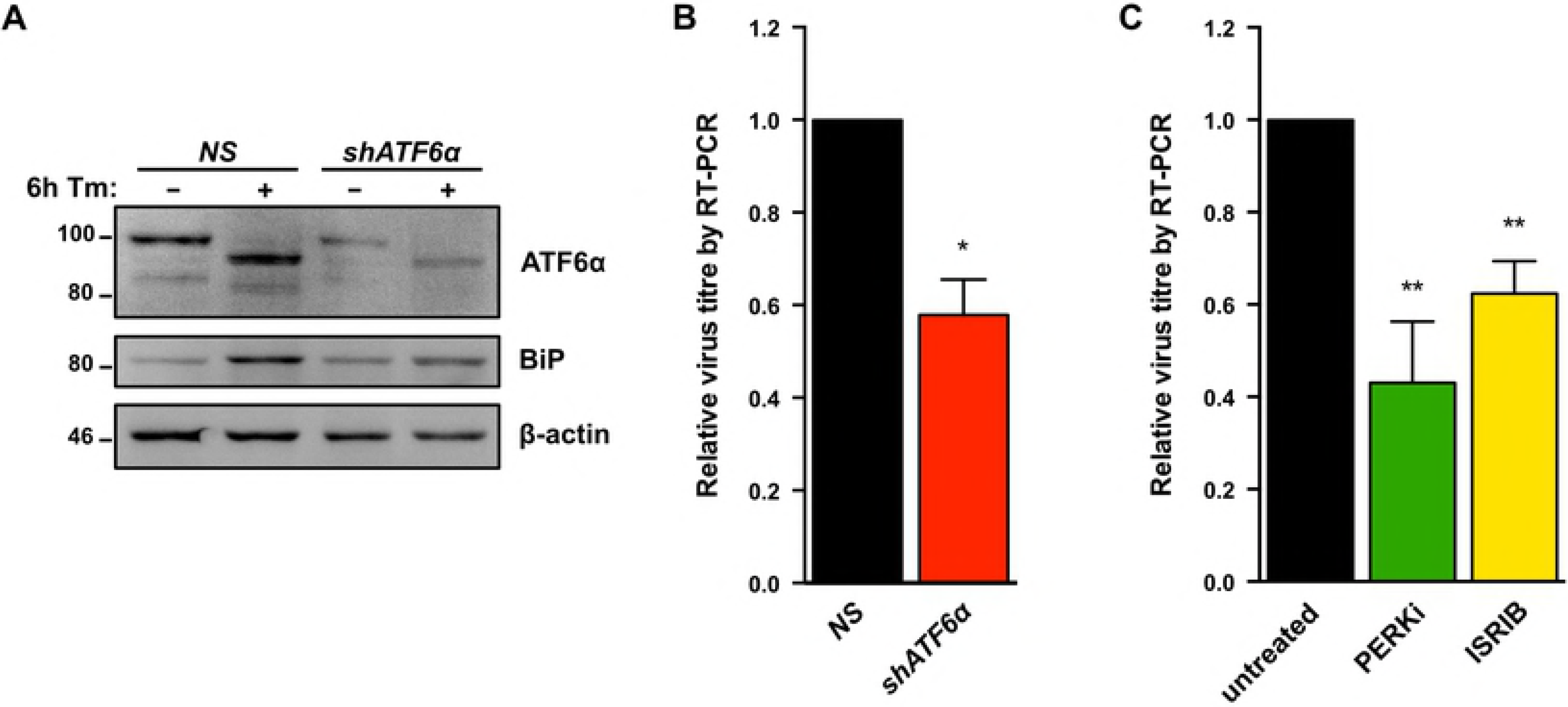
ATF6α and PERK support robust KSHV replication. (A) TREx BCBL1-RTA cells were transduced with lentivirus expressing either non-targeting shRNA (NS) or shRNA against ATF6 and selected for with 1μg/ml puromycin. Following selection, cells were treated with 5ug/ml Tm for 6h and whole cell lysates were analyzed by immunoblots for ATF6 and BiP (Grp78), a transcriptional target of ATF6. (B) As in (A), cells were treated with 1μg/ml doxycycline for 48 hours and virus-protected genomic DNA from cell supernatants were column purified for determining virus titre by qPCR using primers against ORF26. Firefly luciferase plasmid DNA was added during DNA purification for normalization using the ΔΔCq method. (B) TREx BCBL1-RTA cells were treated with 1μg/ml doxycycline for 48 hours with or without 500 nM PERKi GSK2606414 or 250 nM ISRIB, and virus titre was measured by qPCR as in (B). Data are represented as 3 (B) or 4 (C) independent experiments and error bars denote SEM. One-way ANOVA and a post-hoc multiple comparisons test were done to determine statistical significance. (*, p value <0.05; **, p value <0.01).

To determine if IRE1 RNase activity is required for efficient viral replication TREx BCBL1-RTA cells were treated with the IRE1 inhibitor 4μ8c [76], which inhibited virus release in a dose-dependent manner (Fig. 6A). We corroborated these findings in the dox-inducible iSLK.219 cell model that produces KSHV virions bearing GFP transgenes [77]; cells were treated with 4μ8c at the time of dox addition and cell supernatants were harvested 96 hours later, serially diluted, and titered on naive monolayers of 293A cells by flow cytometry. As in the TREx BCBL1-RTA cell model, higher doses of 4μ8c inhibited virion production from iSLK.219 cells, with statistically significant inhibition achieved at the 25 μM dose (Fig. 6B). To confirm these observations, we inhibited IRE1α expression in iSLK.219 cells via RNA silencing. Cells transduced with IRE1α-targeting shRNAs or non-targeting controls were treated with dox for 96 hours, and cell supernatants were once again collected to titer GFP-expressing KSHV virions by flow cytometry. iSLK.219 cells bearing IRE1α shRNAs inhibited release of infectious virions by more than two-fold (Fig. 6C, 6D). Taken together, these data suggest that all three sensors of the UPR are important for robust virus replication.

**Fig. 6:**
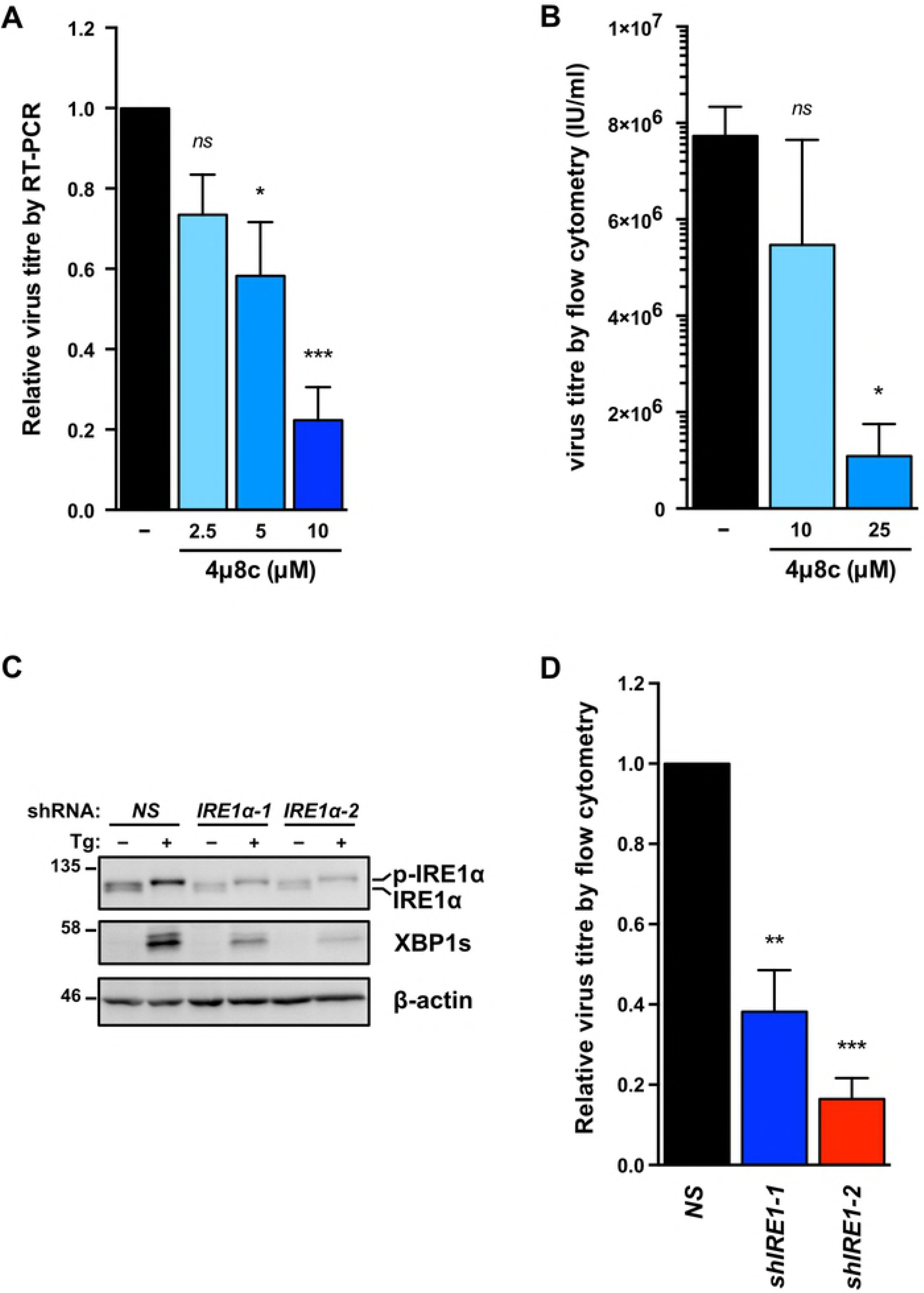
IRE1 is required for efficient KSHV replication. (A) TREx BCBL1-RTA cells were treated with 1μg/ml doxycycline for 48 hours with or without increasing concentrations of IRE1 inhibitor 4μ8c and virus-protected genomic DNA from cell supernatants was measured by qPCR to determine changes in virus titre using the ΔΔCq method by normalizing to *luc2* DNA levels. The mean of four independent experiments are graphed −/+ SEM. (B) iSLK.219 cells were treated with 1μg/ml doxycycline with or without 10 or 25 μM 4μ8c for 96 hours and virus-containing supernatants were serially diluted and spinfected onto a monolayer of 293A cells. GFP-positive cells (infected) were quantified by flow cytometry the following day and virus titre (IU/ml) was calculated as described in the Materials & Methods. The mean of four independent experiments are graphed −/+ SEM. (C) iSLK.219 cells were transduced with two different pLKO.1-blast shRNA lentiviruses targeting IRE1α or an nontargeting control and selected for with blasticidin. Following selection, cells were treated with 150nM Tg for 4 hours and harvested for immunoblot analysis to confirm IRE1α knockdown. The immunoblot shown is representative of two independent experiments performed. (D) Lentivirus transduced iSLK.219 cells from (C), were treated with doxycycline for 96h and virus titre was determined by flow cytometry as previously described. The data are represented as the change in virus titre relative to the non-targeting shRNA control sample and the mean of three independent experiments are graphed −/+ SEM. One-way ANOVA and multiple comparisons test were done to determine statistical significance in (A), (B), and (D). (*ns*, not statistically significant; *, *p* value < 0.05; **, *p* value < 0.01; ***, *p* value < 0.001)

### Ectopic XBP1s expression inhibits release of infectious KSHV virions in a dose-dependent manner

Our studies to this point suggested that efficient lytic replication depends on activation of all three UPR sensor proteins, but the cell fails to produce UPR transcription factors and downstream transcriptional responses required to mitigate ER stress. This suggests that KSHV may re-dedicate the UPR sensors for a new purpose other than to resolve ER stress, and that sustained ER stress may have little impact on viral replication. We also found it puzzling that IRE1 RNase activity was required for efficient lytic replication but XBP1s could not transactivate XBP1s-target genes, including RTA. Furthermore, XBP1s protein levels were reduced following treatment with chemical inducers of ER stress during the lytic cycle despite clear IRE1 activation and efficient *Xbp1* mRNA splicing. For these reasons, we hypothesized that XBP1s accumulation may negatively impact KSHV replication. To complement the XBP1s deficiency in the KSHV lytic cycle we overexpressed XBP1s using dox-inducible lentiviral vectors. We engineered myc-tagged XBP1s to be weakly expressed under the control of a single tet operator (TetO) element, or strongly expressed under the control of seven tandem TetO elements. We transduced iSLK.219 cells with these constructs or an empty vector control, and selected stable cells with blasticidin antibiotic. With the addition of dox, XBP1s and RTA were concurrently expressed from dox-inducible promoters. To demonstrate that the ectopic XBP1s was functional we measured mRNA levels of the XBP1s-target gene *ERdj4* and observed that it is upregulated in cells expressing XBP1s from the 7xTetO compared to empty vector control and peaks at 24 hours post dox addition (Fig. 7A). These cells also displayed higher levels of mRNA and protein for RTA and the RTA target gene ORF45 by 24 hours post-dox treatment; and by 48 hours, mRNA encoding the late viral protein K8.1 was markedly increased compared to controls (Fig. 7A and B), suggesting accelerated viral genome replication. Indeed, intracellular levels of viral genomes at 96 h post-dox were two-fold higher in XBP1s-overexpressing cells compared to empty vector (Fig. 7C). We harvested cell supernatants at 48, 72, and 96 hours post dox and measured virion titer as previously described. Surprisingly, despite accelerated viral gene expression and genome replication in XBP1s-overexpressing cells, there was a dramatic, dose-dependent reduction in virion production by 72 and 96 hours post-dox compared to empty vector control (Fig. 7D). There was also a corresponding 20-fold decrease in release of viral particles by XBP1s-overexpressing cells compared to controls, as measured by qPCR for capsid-protected viral genomic DNA (Fig. 7E). Thus, while ectopic XBP1s expression promotes KSHV lytic gene expression and genome replication, it prevents efficient release of infectious progeny.

**Fig. 7:**
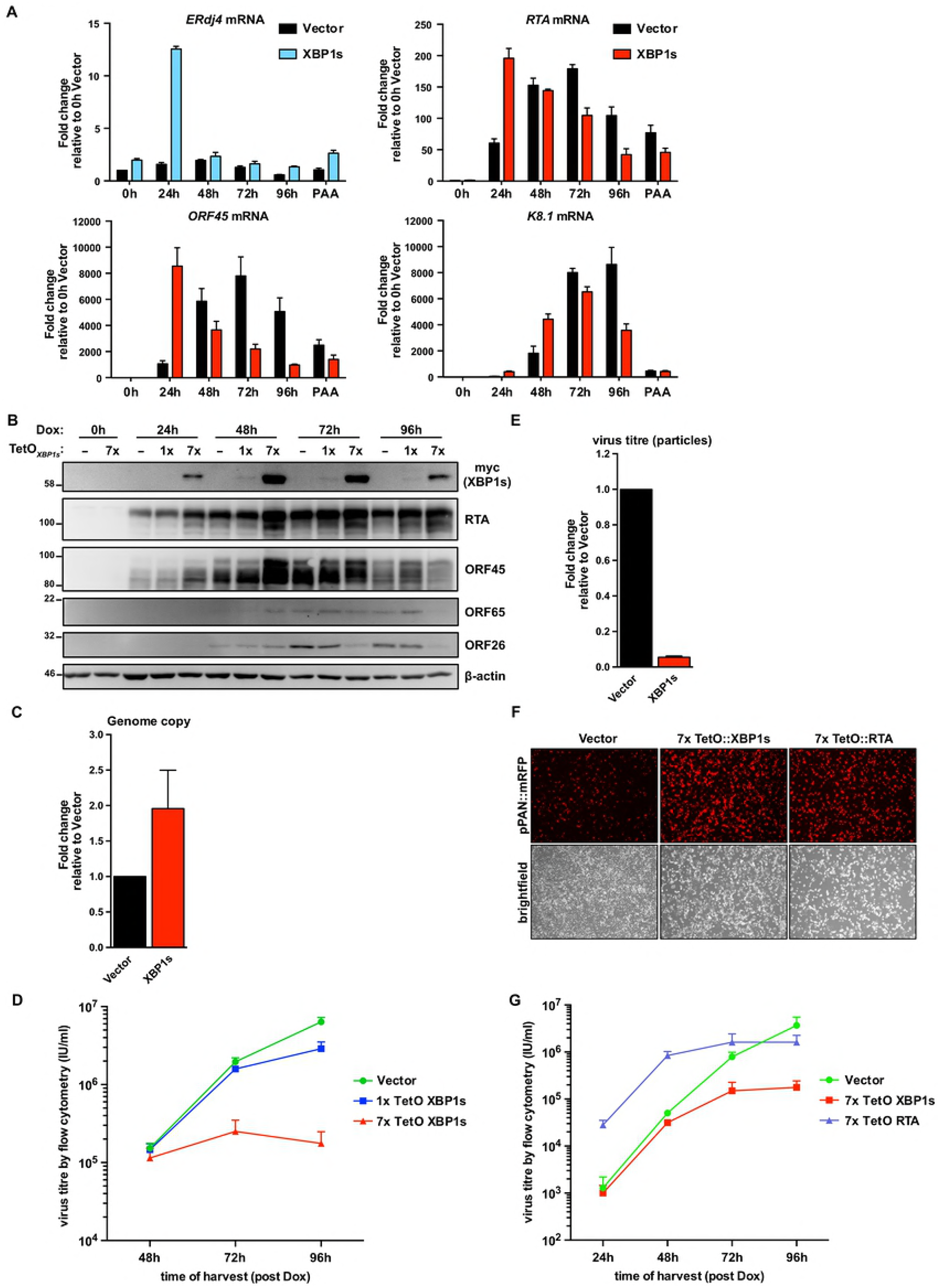
XBP1s overexpression inhibits KSHV replication at a late stage. (A) iSLK.219 cells were transduced with lentiviral doxycycline-inducible expression vectors encoding myc-XBP1s whose expression is driven from 7x tandem tet operator (TetO) for robust gene expression. Following blasticidin expression, lytic replication was induced with doxycycline for 0, 24, 48, 72, and 96 hours or 96 hours with 500 nM PAA (to inhibit genome replication) and and harvested for total RNA. mRNA levels of XBP1s-target gene ERdj4 and viral genes RTA (immediate early gene [IE]), ORF45 (early gene [E]), and K8.1 (late gene [L]) were measured by qPCR using 18S rRNA as a reference gene for normalization. An average of 3 independent experiments are graphed and error bars denote SEM. (B) iSLK.219 cells were transduced with lentiviral doxycycline-inducible expression vectors encoding myc-XBP1s whose expression is driven from either 1x TetO (weak expression) or 7x TetO (strong expression) and treated with doxycycline for the indicated times and harvested for total cell lysates. Immunoblots were done for myc-epitope tag (XBP1s), and the viral proteins RTA (IE), ORF45 (E), ORF65 (L), and ORF26 (L). β-actin was used as a loading control. The presented immunoblots are representative of 2 independent experiments. (C) 7xTetO-myc-XBP1s and vector transduced iSLK.219 cells were treated with doxycycline for 96 hours and intracellular DNA purified. qPCR against ORF26 DNA was done to measure the relative change in viral genome replication using the ΔΔ*Cq* method and normalized to β-actin DNA. Values are the mean of 3 independent experiments −/+ SEM. (D) Virus-containing supernatants from (C) were serially diluted and spinfected onto a monolayer of 293A cells. GFP-positive cells were quantified by flow cytometry the following day and used to calculate virus titre (IU/ml). The values are the mean virus titre of 4 independent experiments −/+ SEM. (E) 7xTetO-myc-XBP1s and vector transduced iSLK.219 cells were treated with doxycycline for 96 hours and DNase-protected genomic DNA from supernatants were column purified. Firefly luciferase plasmid DNA was added during DNA purification to allow for normalization. The relative change in virus titre (infectious and non-infectious) was quantified by qPCR using ORF26 primers and normalized to luciferase DNA using the ΔΔ*Cq* method. Values are the mean of 4 independent experiments −/+ SEM. (F and G) iSLK.219 cells were transduced with lentiviral expression vectors encoding either doxycycline-inducible 7xTetO-myc-XBP1s or 7xTetO-FLAG-RTA. Following blasticidin selection, lytic replication was induced with doxycycline for (F) 48 hours and fluorescence microscopy was used to image RFP-positive cells (cells undergoing lytic replication) or for (G) 24, 48, 72, and 96 hours and virus supernatants were harvested to measure titre by flow cytometry. Values are an average of 3 independent experiments −/+ SEM.

Since XBP1s transactivates the *RTA* promoter, we hypothesized that this defect in virion production could be a negative consequence of *RTA* hyper-activation. We observed a decrease in the accumulation of capsid proteins ORF26 and ORF65 (Fig. 7B), which we speculate could result from RTA hyper-activation and negatively impact late stages of replication [78]. However, experiments with the weaker 1xTetO-XBP1s construct revealed a 2-fold decrease in virion release at 96 hours post dox (Fig. 7D) without affecting RTA, ORF26 and ORF65 protein accumulation (Fig. 7B). To confirm that the diminished production of infectious virions from XBP1s-overexpressing cells is not due to enhanced RTA expression, we also overexpressed RTA in parallel from a 7xTetO in iSLK.219 cells, such that dox addition causes RTA expression by two dox-responsive promoters. KSHV from iSLK.219 cells also express monomeric red fluorescent protein (mRFP) from the viral lytic PAN promoter and can be used to monitor virus reactivation [79]. 48 hours post-dox addition, the levels of mRFP were similar between XBP1s and RTA-expressing cells and noticeably greater than that of the empty vector control (Fig. 7F). We harvested virus-containing supernatants at 24, 48, 72, and 96 hours post-dox and measured virion release by flow cytometry following infection of a naïve 293A monolayer (Fig. 7G). At 24h, when virion production is negligible in empty vector-and XBP1s-expressing cells, there is a significantly higher level of virions produced by 7xTetO RTA-expressing cells which continues up until 72 hours. At 96 hours post-dox, virion production from the 7xTetO RTA-expressing cells hits a plateau. Here again, in agreement with our previous observations, virion production from 7xTetO XBP1s-expressing cells is comparable to empty vector control after 48 hours but by 96 hours post-dox virion production is ~20-fold lower. These data demonstrate that the dramatic reduction in virions produced by XBP1-overexpressing cells is not due to hyperactivation of RTA.

Our findings indicate that XBP1s is both pro-lytic (by transactivating RTA and promoting reactivation from latency) and anti-viral (by inhibiting a late stage in the lytic replication cycle). These confounding observations prompted us to take a step back and corroborate these observations in a different system, wherein XBP1s is expressed from a constitutive CMV promoter in the absence of doxycycline-induced RTA. To achieve this, we transduced iSLK.219 cells with increasing concentrations of lentiviral vectors expressing either myc-tagged XBP1s or FLAG-tagged RTA. To ensure that there were would be an equivalent level of reactivation between cells constitutively expressing RTA or XBP1s, we chose virus dilutions that resulted in a similar range of mRFP levels (indicating lytic cycle initiation) as monitored by fluorescence microscopy (Fig. 8A). Six days after cell transduction, virus-containing supernatants were harvested, and infectious virions were enumerated by infecting naive 293A cells and detecting GFP positive cells by flow cytometry; as a positive control, we harvested virus from untransduced iSLK.219 cells treated with doxycycline for 72h (Fig. 8B). Interestingly, although there is a dose-dependent increase in virion release from RTA-transduced iSLK.219 cells, increasing levels of XBP1s had no impact on virion production and the highest yield of virions was almost 100-fold lower than the corresponding highest yield from RTA-transduced cells or dox-treated iSLK.219 cells. Interestingly, although in RTA-transduced cells, RTA and ORF45 protein levels correspond to the increasing levels of mRFP this does not occur in XBP1-expressing cells; increasing XBP1s expression had a nominal impact on RTA and ORF45 protein accumulation (Fig. 8C). In XBP1s-transduced cells there was a striking reduction in ORF65 accumulation, and ORF45 displayed a mobility shift compared to RTA-transduced cells. Taken together, our observations from multiple systems suggest that XBP1s negatively impacts late stages of KSHV replication resulting in diminished virion yield.

**Fig. 8:**
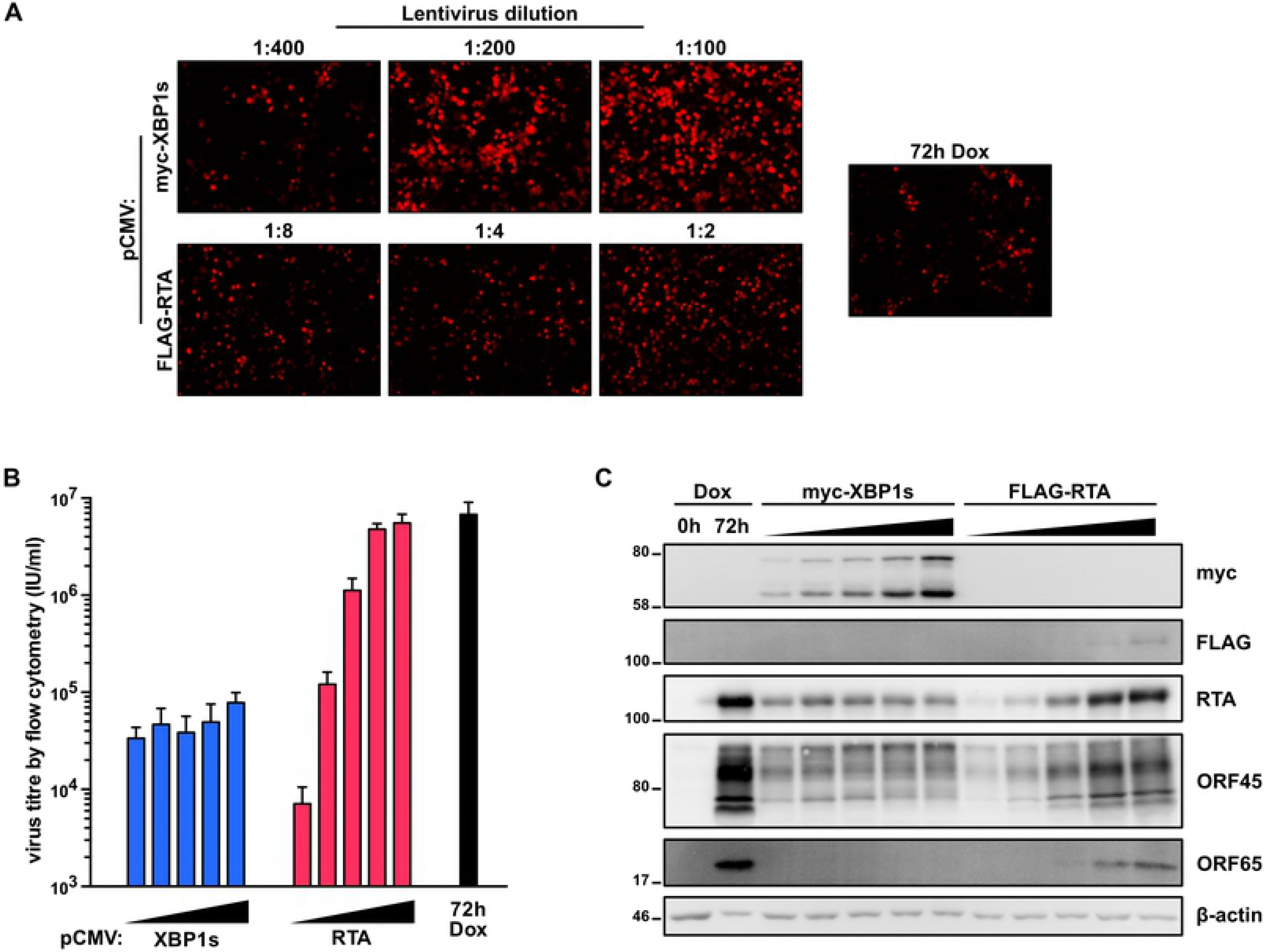
XBP1s overexpression inhibits KSHV replication at a late stage. (A,B, and C) iSLK.219 cells were transduced with increasing concentrations of CMV-driven lentiviral expression vectors encoding myc-XBP1s (LV dilutions: 1:400, 1:200, 1:100, 1:50, 1:25) or FLAG-RTA (LV dilutions: 1:32, 1:16, 1:8, 1:4, 1:2) for 6 days or treated with doxycycline for 72 hours (as a positive control) and (A) a subset of the lentiviral dilutions, cells were imaged for RFP-positive cells by fluorescence microscopy; (B) virus-containing supernatants were harvested for measuring virus titre by flow cytometry; and (C) cell lysates were analyzed by immunoblots for myc-epitope tag (XBP1s), and the viral proteins RTA, ORF45, and ORF65.). β-actin was used as a loading control.

## Discussion

XBP1s-mediated transactivation of the RTA promoter causes reactivation from latency, but little is known about how ER stress and the UPR affect the lytic replication cycle that follows. Here we report that activation of all three UPR sensors (PERK, IRE1, ATF6) is required for efficient KSHV lytic replication, because genetic or pharmacologic inhibition of each UPR sensor diminishes virion production. Despite strong UPR sensor activation during KSHV lytic replication, downstream UPR transcriptional responses were restricted. Paradoxically we found that although XBP1s can trigger reactivation by upregulating RTA, forced expression of XBP1s blocks virion production, despite accelerating lytic gene expression and genome replication. These findings suggest that XBP1s has antiviral properties that KSHV must circumvent during the lytic cycle to ensure efficient viral replication. This provides a tidy rationale for viral blockade of XBP1s production during the lytic cycle, but significantly more work will be required to elucidate precise mechanisms of viral activation of UPR sensors and inhibition of downstream UPR transcriptional responses.

All three UPR sensors are activated concurrently during the lytic cycle, which suggests that they may be responding to canonical protein misfolding events in the ER. Such misfolding could be the result of viral glycoprotein biogenesis, but we think this scenario is unlikely because UPR sensor activation in the BCBL-1 and iSLK.219 cell models precedes viral genome replication and bulk synthesis of most structural viral glycoproteins. We favour an alternative hypothesis that lytic replication disrupts ER chaperone function in some way, perhaps through the direct action of viral proteins or indirect activation of signal transduction pathways. It seems counterintuitive that an enveloped virus would actively trigger ER stress, but we speculate that UPR activation in the earliest stages of lytic replication could help prime the ER for the impending increase in viral glycoprotein synthesis. A more comprehensive accounting of changes in gene expression during early lytic replication is required to determine whether there is indeed an activation of an adaptive UPR to prepare the cell for late gene expression. There are multiple examples of viruses that trigger Ca^2+^ release from ER stores, which may impede protein folding and activate an ER stress response [80]. Indeed, KSHV encodes a viral G protein-coupled receptor (vGPCR) that activates stress signalling and increases cytosolic calcium at least in part by inhibiting the SERCA pump [81]. Thus, vGPCR may act similarly to Tg, inducing broad activation of all three UPR sensors by depleting ER calcium stores essential for protein folding.

Why would a virus cause the induction of ER stress but block the downstream UPR signalling and transcription events that mitigate stress? UPR signalling is etiologically-linked to inflammatory diseases including Crohn’s Disease and type 2 diabetes [34,82,83]. Furthermore, maximal IFN production following Toll-like receptor (TLR) engagement has been shown to require UPR activation and XBP1s activity [84]. Thus, KSHV suppression of UPR transcription may dampen antiviral inflammatory responses. Furthermore, because UPR transcription factors transactivate genes involved in catabolism, we speculate that blockade of their activity may allow newly-synthesized viral gene products to evade degradation. For example, ATF6 and XBP1s transactivate genes involved in ERAD [28,62], so suppression of ATF6/XBP1s transcription could prevent ERAD-mediated degradation of newly-synthesized viral glycoproteins. Similarly, because ATF4 and its downstream target CHOP transactivate autophagy [16] and autophagy restricts herpesvirus replication [85,86], suppression of ATF4 and CHOP accumulation may allow KSHV to evade negative catabolic effects of autophagy. Viral suppression of CHOP synthesis may also prevent the pro-apoptotic effects of CHOP [87], thereby extending the survival of cells enduring the later stages of lytic replication. However, the consequences of allowing unchecked accumulation of misfolded proteins are unclear. Autophagy has been shown to compensate and assist with bulk protein degradation when ERAD is defective or inadequate for degrading misfolded proteins like aggregation-prone membrane proteins [88,89]. Since ERAD genes are not upregulated during lytic replication, it is possible that autophagy is required to clear misfolded proteins. However, there are multiple KSHV proteins that have been shown to directly inhibit autophagy including vFLIP, K-survivin, and vBCL2 [90–92], as well as multiple KSHV proteins that stimulate the pro-growth PI3K-Akt-mTOR signalling pathway [93,94]. Simultaneous inhibition of ERAD and autophagy likely would cause the accumulation of the misfolded proteins, including aggregated membrane proteins, in KSHV infected cells. Mechanisms that KSHV employs (if any) to circumvent a possible accrual of misfolded proteins remain unknown.

How might KSHV inhibit ATF4 protein accumulation? ATF4 synthesis relies on eIF2α phosphorylation, which binds and inhibits the eIF4B guanine nucleotide exchange factor and depletes the available pool eIF2α-tRNA_Met_^i^-GTP ternary complexes; this causes bypass of AUG start sites that normally initiate uORF translation and favours initiation at the *bona fide* downstream ATF4 AUG start codon [14]. Host shutoff during lytic replication decreases the pool of translation-ready mRNAs [95], which may necessitate increased phosphorylation of eIF2α to suppress eIF4B and allow uORF skipping for ATF4 synthesis. ATF4 accumulation is also controlled by post-transcriptional modifications; when the ISR is activated, host methyltransferases add N^6^-methyladenosine (m6A) to the ATF4 mRNA, which further enhances ATF4 ORF translation [96]. The m6A machinery is also required for efficient KSHV lytic replication [97]. Intriguingly, these studies show that knockdown of the m6A methyltransferase METTL3 increases ATF4 synthesis and decreases virion production. Thus, it is possible that KSHV hijacks the m6A machinery to prevent translation of ISR mRNAs while promoting efficient translation of KSHV gene products.

Despite efficient XBP1 mRNA splicing, XBP1s protein failed to accumulate in lytic cells in iSLK.219 and BCBL-1 cell models, even after extended Tg or Tm treatments. We don’t yet understand the underlying mechanism of viral blockade of XBP1s accumulation, but the interconnected nature of the different arms of the UPR will inform our ongoing studies. For example, we have observed diminished levels of total IRE1 protein at later stages of lytic replication. ATF4 and ATF6(N) have both been shown to upregulate IRE1 [24,98]. Thus, during KSHV lytic replication, failure to accumulate the UPR transcription factors may diminish IRE1 protein levels required to support XBP1s synthesis. We are actively investigating these UPR regulatory mechanisms in our KSHV infection models.

Many viruses modulate the UPR, including other herpesviruses. Herpes simplex virus type 1 (HSV-1) has been shown to dysregulate IRE1 activity [99]; HSV-1 infected cells displayed increased IRE1 protein levels but diminished IRE1 RNase activity, and treatment with the IRE1 inhibitor STF-083010 [100] diminished viral replication [101]. Moreover, HSV-1 glycoprotein B (gB) binds and inhibits PERK to support robust viral protein synthesis [57]. Interestingly, ectopic expression of FLAG-tagged XBP1s inhibits HSV-1 replication [101], consistent with our findings in KSHV infection models. By contrast, human cytomegalovirus (HCMV) lytic replication in fibroblast cells selectively activates PERK and IRE1, but not ATF6 [55]. Consistent with our model, IRE1 activation causes normal *Xbp1* mRNA splicing but fails to upregulate the XBP1s target gene EDEM (XBP1s protein levels were not evaluated in this study). However, PERK causes ISR activation and normal ATF4 accumulation during HCMV lytic replication, in sharp contrast to the situation in KSHV infection [55]. A recent study linked HCMV PERK and IRE1 activation to the UL148 glycoprotein [56]. In cells infected with a UL148-deficient HCMV, XBP1s protein levels were diminished compared to wildtype but levels of the XBP1s-target gene EDEM1 are unaffected, suggesting that XBP1s activity may also be inhibited in CMV infected cells. Another study showed that HCMV UL38 induces eIF2α phosphorylation and ATF4 synthesis and promotes cell survival and virus replication during drug-induced ER stress by repressing JNK phosphorylation [102]. HCMV UL50 protein and the murine cytomegalovirus (MCMV) ortholog M50 bind IRE1 and promote its degradation, thereby inhibiting *Xbp1* mRNA splicing [103]. The accumulating evidence indicates that herpesviruses have evolved distinct mechanisms to regulate the UPR to promote viral replication. We suspect that many more viral regulators of the UPR will be discovered in the coming years. Such studies will benefit from the development of appropriate platforms to screen hundreds of viral ORFs for UPR and ISR-modulating activity.

IRE1 is modulated by diverse viruses. Hepatitis C virus (HCV) replication triggers chronic ER stress and sustained activation of all three branches of the UPR [104], but other studies showed that HCV suppresses the IRE1/XBP1 pathway and virus replication is enhanced in IRE1 KO MEFs [105]. Influenza A virus (IAV) specifically activates IRE1 with little or no activation of PERK or ATF4 and chemical inhibition of IRE1 diminishes virion production [106]. Why does this array of unrelated viruses target IRE1? Beyond the aforementioned XBP1s transcription program, IRE1 can activate the JNK signalling pathway [107], which plays important roles in viral replication, cellular stress responses, and cell fate [108,109]. For HSV-1, IRE1 activation could activate JNK, which has been shown to support viral replication in multiple cell types [101]. JNK has also been shown to play important roles in KSHV infection; *de novo* infection triggers JNK activation, and ER stress-induced JNK activation can reactivate KSHV from latency [110,111]. Later stages of KSHV lytic replication also feature JNK activation, and ectopic expression experiments have shown that KSHV lytic proteins K15 and vGPCR can induce JNK phosphorylation [112,113]. It will be interesting to determine whether IRE1 is required for JNK activation during KSHV infection.

IRE1 can also cleave a subset of ER-targeted mRNAs for degradation in a process called RIDD [30]. It is possible that certain viruses may hijack RIDD as a form of host shutoff to ensure preferential translation of secreted viral proteins. Conversely, if viral mRNAs contain XBP1-like cleavage sites, then the virus would likely want to suppress IRE1 RNase activity [114]. The molecular events that direct IRE1 toward RIDD or *Xbp1* mRNA splicing are not well understood but one study suggests that the higher order multimerization of IRE1 can dictate the response, whereby oligomeric IRE1 prefers *Xbp1* as a substrate, whereas dimeric IRE1 preferentially cleaves mRNAs through RIDD [115]. Since the levels of IRE1 are impacted by many of these viruses, including CMV, HSV-1, and KSHV, it will be interesting to determine if IRE1 specificity of mRNAs is impacted during infection. A search for XBP1-like stem loops in herpesviruses mRNAs could help inform if RIDD is antiviral.

XBP1s is required for the differentiation of B cells into antibody-secreting plasma cells [33]. KSHV co-opts the B lymphocyte differentiation program when it infects immature B cells and drives them to differentiate into plasmablast cells [43], which serves as the long-term latent reservoir in the body. It would be interesting to test if the KSHV modulation of the IRE1/XBP1s pathway that we report is required to drive this unique differentiation program. Consistent with our findings and unlike some XBP1s-positive plasmablastic lymphomas, patient-derived PEL cells can lack XBP1s expression [116]. Suppression of XBP1s accumulation may allow KSHV to prevent reactivation from latency and preserve this viral reservoir. If this hypothesis is correct, then ectopic expression of XBP1s should kill KSHV-positive PEL cells, while interfering with viral replication. It is also possible that the lack of XBP1s-positive PEL cells is because XBP1s-mediated plasma cell differentiation should also trigger reactivation and ultimately cell death. The XBP1s transcriptional program during plasma cell development is likely to have distinct players and analysing some of these downstream effectors for possible KSHV inhibition may help identify why XBP1s potently inhibits virus production in our model.

## Materials and Methods

### Cell culture and chemicals

293A (ThermoFisher), HEK293T (ATCC), Hela Tet-Off (Clontech), and iSLK.219 cells (a kind gift from Don Ganem) were cultured in DMEM (ThermoFisher) supplemented with 10% heat-inactivated fetal bovine serum (FBS), 100 Units/mL penicillin, 100 μg/mL streptomycin, and 2mM L-Glutamine. iSLK.219 cells were also passaged in the presence of 10 μg/mL of puromycin (ThermoFisher) to maintain the rKSHV.219 episomal DNA. TREx BCBL1-RTA cells (a kind gift from Jae Jung) were cultured in RPMI-1640 supplemented with 10% heat-inactivated FBS, 500 μM β-mercaptoethanol and the same concentrations of penicillin, streptomycin, and L-Glutamine as the adherent cell lines. All cells were maintained at 37°C with 5% CO_2_.

To induce lytic replication via expression of the RTA transgene in TREx BCBL1-RTA and iSLK.219 cells, 1 μg/mL of doxycycline (dox; Sigma) was added to the cells. iSLK.219 cells were seeded at density of 2×10^5^ cells/well of a 6-well plate one day prior to lytic induction and TREx BCBL1-RTA cells were seeded at a concentration of 2.5×10^5^ cells/mL immediately prior to lytic induction.

4μ8c (Axon Medchem and Sigma), PERKi (GSK2606414; Tocris), ISRIB (Sigma), Thapsigargin (Sigma), and Tunicamycin (Sigma) were dissolved in DMSO (Sigma) to stock concentrations and diluted to the indicated concentrations in cell media.

### Plasmids

To generate the plasmids pLJM1 B* Puro and pLJM1 B* BSD, the genes encoding puromycin N-acetyltransferase and blasticidin S deaminase were amplified from pBMN-IRES-Puro and pBMN-IRES-Blast, respectively (previously generated in the lab by switching GFP with PuroR or BlastR from pBMN-I-GFP that was generated by the Nolan Lab [Stanford]), with forward and reverse primers containing BglII and KpnI RE sites. The antibiotic selection cassettes were cut-and-pasted into pLJM1.D (previously generated in the lab from modifying the MCS of pLJM1 plasmid that was generated by the Sabatini Lab [MIT]), deleting the BamHI RE site. A new MCS was generated by annealing overlapping oligos containing NheI and MfeI forward and reverse RE sites and inserted into the plasmid digested with NheI and EcoRI to replace the existing MCS with unique RE sites in the following order: NheI, AgeI, BamHI, EcoRI, PstI, XbaI, MluI, SalI, EcoRV. 1x TO (tet operator) and 7x TO promoters were generated by annealing overlapping oligos containing one or seven copies of the tetracycline operator (5’- TCCCTATCAGTGATAGAGA-3’) and a minimal CMV promoter and using PCR extension to amplify a blunt oligo containing NdeI and NheI RE sites. The amplicon was digested with NdeI and NheI and pasted into the pLJM1 B* BSD replacing the CMV promoter.

To generate pLJM1 B* Puro HA-ATF6, HA-ATF6 was PCR amplified from pCGN-ATF6 (a gift from Ron Prywes, Addgene plasmid # 11974) and cut and pasted into pLJM1 B* Puro with NheI and AgeI. pCMV2B-XBP1s was generated by PCR amplifying XBP1 from total RNA isolated from TREx BCBL1-RTA cells treated with Tg for 4h. After PCR, DNA was digested with PstI (cleaves unspliced XBP1 DNA only) to enrich for the spliced isoform of XBP1 and cut and pasted into pCMV2B with BamHI and XhoI and in-frame of the N-terminal FLAG-tag. pLJM1 BSD CMV, 1x TO, and 7x TO myc-XBP1s were generated by and including the myc-tag nucleotide sequence in the 5’ primer and amplifying XBP1s from pCMV2B-XBP1s. Cloned myc-XBP1s was subsequently cut and pasted into their corresponding lentiviral plasmids with BamHI and SalI. FLAG-RTA was cut from pcDNA3-FLAG-RTA (previously generated in the lab) with RE sites EcoRI and XhoI and pasted into pLJM1 B* BSD CMV and 7x TO with RE sites EcoRI and SalI (via compatible sticky ends between XhoI and SalI).

### Lentiviral vectors

HEK293T cells were seeded on 10cm plates for 60-70% cell density the following day. Cells were transfected with polyethylenimine (PEI; Sigma) and the following plasmids for lentiviral generation: pLJM1 transfer plasmid, pMD2.G, and psPAX2. pMD2.G and psPAX2 are gifts from Didier Trono (Addgene plasmids # 12259 and # 12260). 48 hours post transfection, lentivirus containing supernatants were passed through a 0.45μM filter and frozen at −80°C. For transducing iSLK.219, iSLK.219 cells were seeded at density of 5×10^4^ cells/well of a 6-well plate and the following day were resuspended in media containing 4 μg/mL polybrene (Sigma). Lentivirus was serially diluted dropwise onto cells and incubated at 37°C for 24 hours. Following infection, the media was refreshed containing 10 μg/mL blasticidin. For transducing TREx BCBL1-RTA cells, cells were seeded at a concentration of 2.5×10^5^ cells/mL in media containing the same concentration of polybrene and lentivirus was serially diluted onto cells. 24 hours post infection, the media was replaced with media containing 1 μg/mL puromycin. The first lentivirus dilution that resulted in minimal cell death following antibiotic selection was used for subsequent experiments.

### Luciferase Assays

4x pORF50XRE-FLuc plasmid was generated by annealing oligos that have the XBP1 response element sequence 5’-ATGACACGTCCC-3’ found in ORF50 promoter (Dalton-Giffin *et al.* 2009 J.Virol) and is repeated 4 times flanked by 10bp repeats. The oligo also contains KpnI and XhoI RE sites and was inserted into the firefly luciferase plasmid pGL4.26[*luc2*/minP/Hygro] (Promega).

Hela tet-off cells were seeded at a density of 3.2×10^4^ cells/well of a 12-well plate and transfected the following day with FuGENE HD transfection reagent and the following plasmids per well: either 400 ng pCR3.1 Vector control or pCMV2B-XBP1s, 80ng 4x pORF50XRE-FLuc, and 20ng of the *Renilla* luciferase plasmid pRL-TK for normalization. 24 hours post transfection, cells transfected with pCR3.1 were untreated or treated with 150 nM Tg for 16h. Following treatment, firefly and *Renilla* luciferase activity was detected by Glomax 20/20 Luminometer (Promega) after incubating with corresponding substrates from the Dual-Luciferase Reporter System (Promega) according to the manufacturer’s instructions. Firefly luciferase relative lights units were normalized to *Renilla* luciferase and expressed as fold change in signal relative to untreated vector control.

### shRNA lentivirus cloning and knockdown

The pLKO.1-TRC control plasmid was a gift from David Root (Addgene plasmid # 10879) and the RNAi Consortium was used to design the following oligos for generating shRNA lentiviral vectors against ATF6α and IRE1α with the targeting sequences underlined (Fwd/Rev): shAFT6α (TRCN0000416318): 5’-CCG<u>GACAGAGTCTCTCAGGTTAAAT</u>CTCGAGATTTAACCTGAGAGACTCTGTTTTTTG/5’-AATTCAAAAAACAGAGTCTCTCAGGTTAAATCTCGA<u>GATTTAACCTGAGAGACTCTGT</u>-3’ shIRE1α-1 (TRCN0000356305): 5’-CCGGTCAACGCTGGATGGAAGTTTGCTCGAGCAAACTTCCATCCAGCGTTGATTTTG -3’/5’-AATTCAAAAATCAACGCTGGATGGAAGTTTGCTCGA<u>GCAAACTTCCATCCAGCGTTGA</u>-3’shIRE1α-2 (TRCN0000235529): 5’-CCG<u>GAGAGGAGGGAATCGTACATTT</u>CTCGAGAAATGTACGATTCCCTCCTCTTTTTTG-3’/5’-AATTCAAAAAAGAGGAGGGAATCGTACATTTCTCGA<u>GAAATGTACGATTCCCTCCTCT</u>-3’ The cloning strategy on Addgene (http://www.addgene.org/tools/protocols/plko/) was used to clone the oligos into either pLKO.1-Puro or pLKO.1-Blast and lentivirus generation was completed as described previously.

### Fluorescent imaging

For Fig.7, iSLK.219 cells were seeded at density of 2×10^5^ cells/well of a 6-well plate and and 48 hours post addition of doxycycline, brightfield images and fluorescent images were captured with Olympus CKX41 microscope fitted with a QImaging QICAM Fast 1394 digital camera and Lumencor Mira light engine and using the 10x objective. For Fig. 8, iSLK.219 cells were seeded at density of 5×10^4^ cells/well of a 6-well plate and transduced with the indicated lentiviral vectors. Fluorescent images were captured with an EVOS FL Cell Imaging System with 10x objective (ThermoFisher) following 5 days post transduction or 48 hours post dox.

### Immunoblotting

Cells were washed once with ice-cold PBS and lysed in 2x Laemmli Buffer (120mM Tris-HCl pH 6.8, 20% glycerol, 4% SDS). Lysates were passed through a 21-gauge needle 5-7 times to minimize viscosity and protein concentration was quantified with DC protein assay (Bio-Rad). DTT and bromophenol blue were added to samples, boiled, and 10-20 μg of protein per sample were loaded on 6-12% polyacrylamide gels and resolved by SDS-PAGE. Protein was transferred to PVDF membranes using the Trans-Blot Turbo Transfer System (Bio-Rad) and blocked in 5% skim milk diluted in Tris-buffered-saline (TBS) supplemented with 0.1% Tween 20 (Fisher Bio) (TBS-T) for 1 hour at room temperature followed by overnight incubation at 4°C with primary antibody diluted in 5% bovine serum albumin in TBS-T. Following washing of primary antibody, membranes were incubated with IgG HRP-linked antibodies followed by exposure to ECL-2 western blotting substrate (Thermo Scientific, Pierce) and imaged on a Carestream Image Station 400mm Pro (Carestream) using the chemifluorescence setting. The following antibodies were used: PERK (Cell Signaling; #5683); IRE1α (Cell Signaling; #3294); ATF6α (Abcam; ab122897); XBP1 (Cell Signaling; #12782); Phospho-eIF2α (Ser51; Abcam; ab32157); Total eIF2α (Cell Signaling; #5324); ATF4 (Santa Cruz; sc-200); Myc-tag (Cell Signaling; #2278); FLAG-tag (DYKDDDDK tag; Cell Signaling; #8146); ORF45 (Thermo-Fisher; MA5-14769); ORF65 (a kind gift from Shou-Jiang Gao); RTA (a kind gift from David Lukac); β-actin HRP conjugate (Cell Signaling; #5125); anti-rabbit IgG HRP-linked (Cell Signaling; #7074); antimouse IgG HRP-linked (Cell Signaling; #7076).

### Xbp1 RT-PCR splicing assay

RNA was isolated from TREx BCBL1-RTA cells with the RNeasy Plus Kit (Qiagen) and 500 ng total RNA was reverse transcribed with qScript cDNA SuperMix (Quanta) according to manufacturers’ protocols. Based on the Ron Lab protocol (http://ron.cimr.cam.ac.uk/protocols/XBP-1.splicing.06.03.15.pdf) and minor modifications, a 473 bp PCR product overlapping the IRE1 splice site was amplified with XBP1 Fwd primer (5’-AAACAGAGTAGCAGCTCAGACTGC-3’) and XBP1 Rev primer (5’-TCCTTCTGGGTAGACCTCTGGGAG-3’). The amplified PCR product was digested overnight with 40 units of High Fidelity PstI (New England Biolabs) to cleave unspliced XBP1 cDNA. The PCR products were resolved on a 2.5% agarose gel made with 1x TAE (Tris-acetate-EDTA) buffer and stained with SYBR Safe (ThermoFisher) and visualized on a ChemiDoc Imaging Station (Bio-Rad). Percent *Xbp1* mRNA splicing was calculated by densitometry analysis with Image Lab software ver. 6.0.0 (Bio-Rad) and calculated using the following formula:

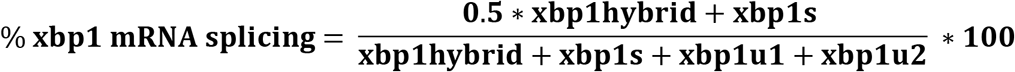

### Quantitative Reverse-Transcription PCR (RT-qPCR)

RNA was isolated from TREx BCBL1-RTA cells with the RNeasy Plus Kit (Qiagen) and 500 ng total RNA was reverse transcribed with qScript cDNA SuperMix (Quanta) according to manufacturers’ protocols. A CFX96 Touch Real-Time PCR Detection System (Bio-Rad) and GoTaq qPCR MasterMix (Promega) was used to perform Real-Time PCR. Changes in mRNA levels were calculated by the ΔΔCq method and normalized using 18S rRNA as a reference gene. The following primer sets were used in the study:
*GRP78* F: 5’-GCCTGTATTTCTAGACCTGCC-3’; R: 5’-TTCATCTTGCCAGCCAGTTG-3’ *HERPUD1* F: 5’-AACGGCATGTTTTGCATCTG-3’; R: 5’-GGGGAAGAAAGGTTCCGAAG-3’ *EDEM1* F: 5’-TTGACAAAGATTCCACCGTCC-3’; R: 5’-TGTGAGCAGAAAGGAGGCTTC-3’ *ERdj4* F: 5’-CGCCAAATCAAGAAGGCCT-3’; R: 5’-CAGCATCCGGGCTCTTATTTT-3’ *ORF26* F: 5’-CAGTTGAGCGTCCCAGATGA-3’; R: 5’-GGAATACCAACAGGAGGCCG-3’ *ORF50* F: 5’-GATTACTGCGACAACGGTGC-3’; R: 5’-TCTGCGACAAAACATGCAGC *K8.1* F: 5’-AGATACGTCTGCCTCTGGGT-3’; R: 5’-AAAGTCACGTGGGAGGTCAC-3’ *ORF45* F: 5’-TGATGAAATCGAGTGGGCGG-3’; R: 5’-CTTAAGCCGCAAAGCAGTGG-3’ *18S* F: 5’-TTCGAACGTCTGCCCTATCAA-3’ R: 5’-GATGTGGTAGCCGTTTCTCAGG-3’

### Viral genome amplification

DNA was harvested from iSLK.219 cells with DNeasy Blood & Tissue Kit (Qiagen) according to manufacturer’s protocol. RT-PCR was carried out as described previously with primers specific to KSHV ORF26 (as listed previously) and β-actin (F: 5’-CTTCCAGCAGATGTGGATCA-3’; R: 5’-AAAGCCATGCCAATCTCATC-3’). Changes in KSHV genome copy number was calculated by the ΔΔCq method and normalized to β-actin.

### Titering DNAse-protected viral genomes

Virus containing cell supernatants were processed by first pelleting floating cells and debris and then 180 μL cleared supernatant were treated with 20 μL of 3 mg/mL DNase I (Sigma) for 30 minutes at 37°C. Viral genomic DNA was then purified from the supernatant with DNeasy Blood & Tissue Kit (Qiagen) according to the manufacturer’s protocol with the following modifications: 10 μg of salmon sperm DNA (Invitrogen) and 1 ng of luciferase plasmid pGL4.26[*luc2*/minP/Hygro] (Promega) were added to the lysis buffer. RT-PCR was performed as described previously with primers specific to KSHV ORF26 (as listed previously) and *luc2* (F: 5’-TTCGGCAACCAGATCATCCC-3’; R: 5’-TGCGCAAGAATAGCTCCTCC-3’). Changes in virus titre was calculated by the ΔΔCq method and normalized to *luc2*

### rKSHV.219 infection and titering

Virus-containing supernatant was harvested from iSLK.219 cells at the indicated times by pelleting cellular debris at 3300 x g for 5 minutes and then stored at −80°C until ready to titer the virus. 1×10^5^ 293A cells/well were seeded in 12-well plates to obtain a confluent monolayer two days later. The thawed viral inoculum was briefly vortexed and centrifuged again at 3300 x g for 5 minutes. Two-fold serial dilutions of viral supernatants were applied to the monolayer containing 4 μg/ml polybrene and 25 mM HEPES (Gibco) and centrifuged at 800 x g for 2 hours at 30°C. The total cell count per well was also determined from an uninfected well. Fresh media was applied immediately after spinoculation. 20-24 hours post infection, two dilutions that resulted in less than 30% GFP-positive cells (the linear range for infection [data not shown]) were trypsinized, washed once with PBS and fixed with 1% paraformaldehyde in PBS. GFP-positivity was measured on either FACSCalibur or FACSCanto cytometers (BD) by gating on FSC/SSC and counting 10000-15000 “live” events. Gating and % GFP positive events was determined with FCS Express 6 Flow Cytometry Software (ver.6.0; De Novo). Virus titre was calculated as IU/mL with the following formula:

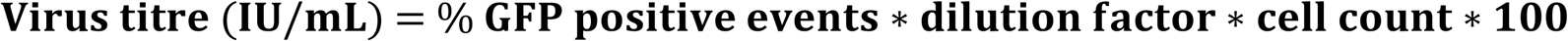

The two dilution series were averaged for the final titre value.

### Statistical analysis

Prism7 (GraphPad) was used for generating graphs and performing statistical analysis. Unpaired Student’s t-tests were used to determine significance between two groups. One-way or two-way ANOVA was used to compare multiple samples or between grouped samples respectively, and an appropriate post-hoc test was done to determine differences between groups. p-values <0.05 were considered significant and denoted as the following: <0.05 (*), <0.01 (**), <0.001 (***), <0.0001 (****)

## Acknowledgements

We thank members of the McCormick lab for critical reading of this manuscript. We would like to thank Don Ganem (Novartis), Shou-Jiang Gao (USC), Jae Jung (USC), and David Lukac (Rutgers) for providing reagents.

## Supplemental Figure. 1: XBP1s transactivates RTA to induce lytic reactivation

(A) Wildtype BCBL1 cells were treated with either 1mM sodium butyrate (NaB) and 20 ng/mL of the phorbol ester TPA (12-O-Tetradecanoylphorbol-13-acetate) or 150 nM Tg for 6, 12, and 24 hours and RTA mRNA levels were measured by qPCR. Changes in mRNA levels were calculated by the ΔΔ*Cq* method and normalized using β-actin. An average of 3 independent experiments are graphed and error bars denote SEM. (B) Hela Tet-Off cells were transfected with pCMV2B-XBP1s or pCR3.1 Vector control and a firefly luciferase expression construct that is driven by 4 tandem XBP1 target sequences that are derived from the promoter of ORF50. A renilla luciferase plasmid was added for normalization and 24 hours later the vector control samples were treated with 500 nM Tg for 16h. Approximately 48 hours post transfection, cells were lysed and a dual-luciferase assay was conducted to measure XBP1 activity.

